# Domestic pigs experimentally infected with *Mycobacterium bovis* and *Mycobacterium tuberculosis* exhibit different disease outcomes

**DOI:** 10.1101/2021.10.25.465759

**Authors:** Nirajan Niroula, Ze Long Lim, Stewart Walker, Yanyun Huang, Volker Gerdts, Slim Zriba, Kylee Drever, Jeffrey M. Chen

**Author notes:** Corresponding Authors: Mailing address: Vaccine and Infectious Disease Organization (VIDO) University of Saskatchewan, 120 Veterinary Road, Saskatoon, SK, S7N 5E3. Canada, and.

## Abstract

Domestic pigs share many similarities with humans in their pulmonary anatomy, physiology, and immunology. Accordingly, pigs have been shown to be valuable models to study human tuberculosis (TB). Here we examined the outcome of disease in domestic pigs challenged via two different routes with either the human-adapted TB bacillus *Mycobacterium tuberculosis* or the zoonotic bovine TB bacillus *M. bovis* in head-to-head comparisons. We found that pigs challenged intravenously with *M. bovis* AF2122/97 exhibited severe morbidity and rapid onset of mortality, accompanied by higher tissue bacterial burden and necrosis compared to pigs challenged similarly with *M. tb* Erdman. Concordantly, pigs challenged with aerosolized *M. bovis* AF2122/97 exhibited reduced weight gain and more severe pathology than pigs challenged similarly with *M. tb* Erdman. Moreover, pigs aerosol-challenged with *M. bovis* AF2122/97 exhibited a spectrum of granulomatous lesions ranging from small well-contained granulomas to caseous-necrotic lesions mimicking active TB disease in humans. In contrast, pigs aerosol-challenged with *M. tb* Erdman exhibited arrested granuloma development. Irrespective of challenge dose and pathological outcome however, peripheral IFN-γ responses were similar in both *M. bovis* AF2122/97 and *M. tb* Erdman challenged pigs. This study demonstrates domestic pigs can support infections with *M. bovis* and *M. tb* and develop pathology similar to what is observed in humans. And although *M. bovis* AF2122/97 appears to be more virulent than *M. tb* Erdman, both strains can be used to model TB in domestic pigs, depending on whether one wishes to recapitulate either acute and active TB or latent TB infections.

## Introduction

Tuberculosis (TB) is a pulmonary infectious disease that affects a wide range of mammals, and it is one of the leading causes of death in humans. In fact, 1.4 million people globally are estimated to have died in 2019 alone due to TB [1]. TB is caused by members of the *Mycobacterium tuberculosis* complex (MTBC). Most cases of human TB are caused by *Mycobacterium tuberculosis* (*M. tb*) although *M. bovis,* which is another member of the MTBC that affects livestock and wildlife, can cause zoonotic TB [1]. TB transmission occurs mainly through bacteria-containing aerosol particles and primarily affects the lungs. Extra-pulmonary TB like meningeal and miliary TB on the other hand, occurs mostly in children and immune-compromised people respectively [2, 3]. TB manifests as two clinical forms - latent infection which is asymptomatic and non-transmissible, and an active disease which is symptomatic as well as transmissible. However, it is now believed that the two forms are the extremes of a dynamic pathological continuum [4].

Much of our current understanding of TB pathogenesis is based on studies of the disease in animal models that include but are not limited to mice, hamsters, guinea pigs, rabbits and non-human primates (NHPs) [5–7]. Except for NHPs however, none of the other models recapitulate the full spectrum of pathology and clinical signs of human TB [6, 8]. For example, *M. tb*-infected mice which are the most frequently used model, tend to produce pseudo-granulomas and do not recapitulate different aspects of human TB pathology [9]. Guinea pigs are hyper-susceptible to *M. tb,* develop disseminated multi-organ infections and rapidly succumb to the disease [10]. Rabbits, unlike mice and guinea pigs, develop cavitary lung lesions upon infection with virulent MTBC strains [11]. However, they do not mimic the dynamic pathological changes seen in humans and the clinical signs are not obvious which limits their usefulness as a model [6, 12].

Non-human primates (NHPs) remain the best models for human TB due to their close phylogenetic relationship with humans [13, 14], and their inherent susceptibility to infection with MTBC [15, 16]. Indeed, a relatively low dose (10 - 100 CFUs) of *M. tb* is sufficient to establish infection in NHPs and induce a wide spectrum of clinical outcomes ranging from no disease to a rapidly progressing fulminant disease to chronic infection [14, 17]. As in humans, TB granulomas in NHPs are well-organized, consisting of a central core of infected and non-infected macrophages surrounded by a layer of lymphocytes and multi-nucleated giant cells enclosed by a fibrous capsule [17, 18]. Likewise, the formation of caseous necrotizing granulomas seen in humans with active TB also occurs in NHPs [19]. Despite these features, the NHP model has its drawbacks. First, it can be challenging to procure NHPs and facilities capable of maintaining large NHP colonies are limited [20]. Second, NHPs require a lot of space, which can be challenging in biocontainment level-3 labs. Finally, NHP use in research is becoming less socially acceptable due to their well-developed emotional and social behaviors [21].

Due to its similarities to humans with respect to anatomy and physiology [22], the domestic pig (*Sus scrofa domesticus*) has become an increasingly attractive model for studying human diseases like pertussis, influenza, genetic disorders like cystic fibrosis, and xenotransplantation [22–24]. Moreover, the immune system of a pig is similar to that of a human by more than 80 % of the studied parameters compared to a mere 10 % similarity between a human and a mouse [25]. For instance, out of 419 clusters of differentiation (CD) molecules found on the cell surface of humans leukocytes, 359 orthologs have been described in pigs [26]. Likewise, both share similarities in T-cells subtypes and Toll-like receptors which are critical for host-pathogen interactions and immune responses [27]. Both human and porcine lungs are divided into different lobes, which are further demarcated into multiple lobules by connective tissue septa, a feature shown to be important for granuloma encapsulation [28, 29]. The larger size of pigs enables repeated sampling, and the availability of porcine specific reagents has also increased. Since pigs are a livestock species and reared commercially in large numbers, there is a steady supply for medical research.

Wild boars (*Sus scrofa*) infected naturally with *M. bovis* show similar pathology to humans infected with *M. tb* [30, 31]. These include a range of well-contained, non-necrotizing granuloma to caseous-liquifying and calcified lesions [31]. Bolin *et. al.* were the first to show experimental disseminated *M. bovis* infection in mixed-breed domestic pigs [32]. However, the route of challenge in their study was either intravenous, intra-tracheal, or tonsillar deposition, none of which mimic the natural route of MTBC infection [32]. Gil *et.al* on the other hand challenged minipigs with *M. tb* and showed the pig’s pulmonary lobular structure helps establish fibrosis and encapsulation to contain the infection [29]. Likewise, Ramos *et.al* used minipigs to study TB infection and natural transmission in adolescents and infants [33]. Although they challenged minipigs with infectious aerosols of the hypervirulent Beijing strain of *M. tb,* they did not compare *bovis* and *M. tb* [33].

In this study, we infected domestic pigs (*Sus scrofa domesticus*) with *M. tb* Erdman and *M*. *bovis* AF2122/97 strains. Although *M. tb* and *M. bovis* have been compared in cattle and goats [34, 35], we wanted to examine differences in outcomes upon infection with these two MTBC members in pigs. In the first trial, pigs were challenged intravenously with *M. tb* and *M. bovis.* In the second trial, pigs were challenged either with *M. bovis* or *M. tb* at two different doses via the aerosol route. We evaluated clinical signs, size and extent of lesion development and histopathology. In both trials, *M. bovis* induced more severe clinical symptoms and pathology and compared to *M. tb*.

## Results and Discussions

### 1. Intravenous (IV) challenge

#### 1.1. Clinical Signs

Pigs challenged intravenously (IV) with *M. bovis* AF2122/97 exhibited signs of elevated respiration and reduced activity by as early as 7 days post-challenge (pc). All six pigs in the *M. bovis* group exhibited abdominal breathing, stooped posture, inappetence and reduced mobility by day 9 pc (Supplementary file 1). One out of six *M. bovis* challenged pigs died on day 11 pc, while two more pigs reached their humane endpoint and had to be euthanized on day 14 pc. By day 16, the remaining *M. bovis* challenged pigs also reached their humane endpoints and were euthanized (Figure 1). In stark contrast, *M. tb* challenged pigs appeared healthy, ate, and behaved normally, and showed no signs of respiratory distress (Supplementary file 2). Five out of six pigs of this treatment group went on to survive the duration of the trial (Figure 1). There was an earlier onset of fever in pigs challenged with *M. bovis* with all six pigs in the group exhibiting increasing body temperatures that started 3 days pc before peaking around 9 to 10 days pc (Figure 2). In contrast, *M. tb* infected pigs exhibited delayed onset of fever with increases in body temperatures that started 6 to 7 days pc before peaking at about 14 days pc (Figure 2). The body weights of pigs were also measured on the day of the challenge and then at multiple time points pc for the duration of the trial. Although most *M. bovis* challenged pigs survived till 16 days pc, there was a clear indication these pigs were losing more weight compared to the *M. tb* challenged group. Indeed, *M. bovis* infected pigs that were still alive weighed 1.5 kg less on average than *M. tb* infected pigs after two weeks despite starting out at similar weights. Although *M. tb* infected pigs did not lose weight for the entire 35 days of the trial, they did not achieve the type of weight gain expected in uninfected pigs over the same period (Figure 3).

**Figure 1.**
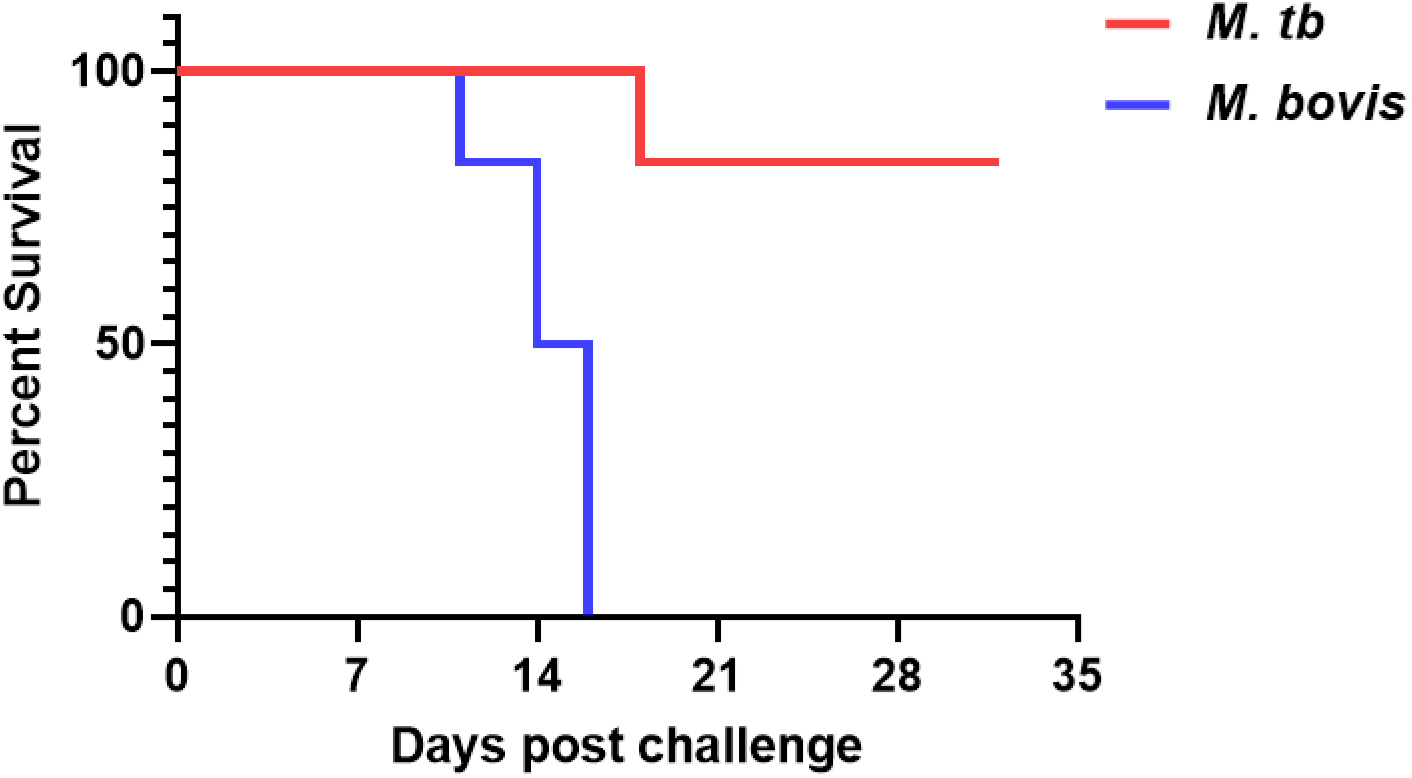
Kaplan-Meier curve of survival rate of pigs challenged intravenously with *M. tb* and *M. bovis*. Pigs challenged with 3 x 10^8^ CFU of *M. bovis* (blue line) died within 16 days post-challenge. In contrast, pigs challenged similarly with *M. tb* showed greater survival and only 1 out of 6 pigs died during the 35-day period (red line). A significant difference was found between *M. tb* and *M. bovis* challenged pigs with respect to the chance of survival post-challenge. (Log-rank test: Chi-square = 10.78, p = 0.001)

**Figure 2.**
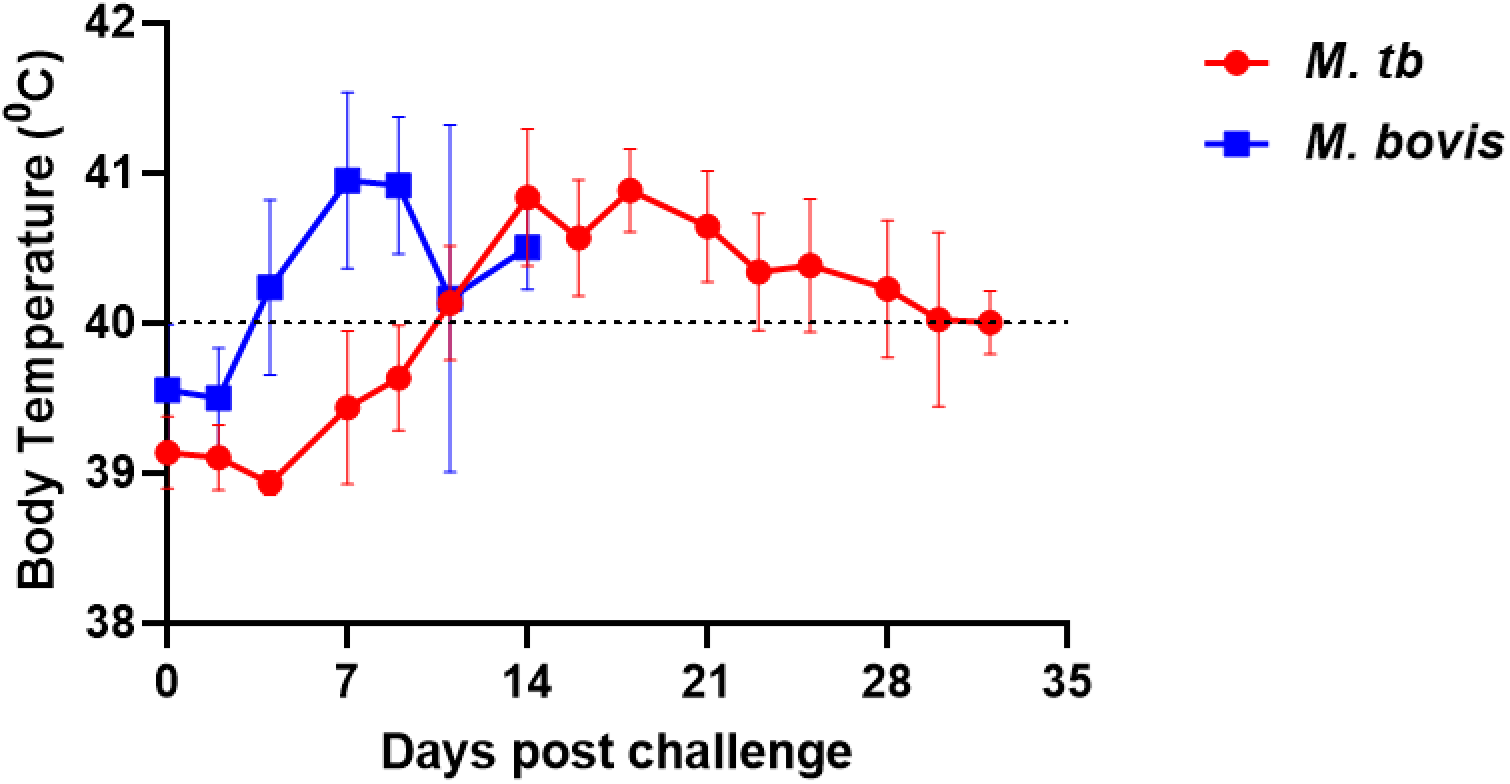
Differences in body temperature (mean ± SD) between *M. tb* and *M. bovis* challenged pigs by intravenous route. Earlier onset of fever was observed in pigs challenged intravenously with *M. bovis* (blue line), which peaked at 7 days post-challenge. Meanwhile, *M. tb* challenged pigs (red line) exhibited delayed onset of fever which peaked around 14 days post-challenge. Temperatures above 40^0^C was considered a sign of fever - indicated by dashed line.

**Figure 3.**
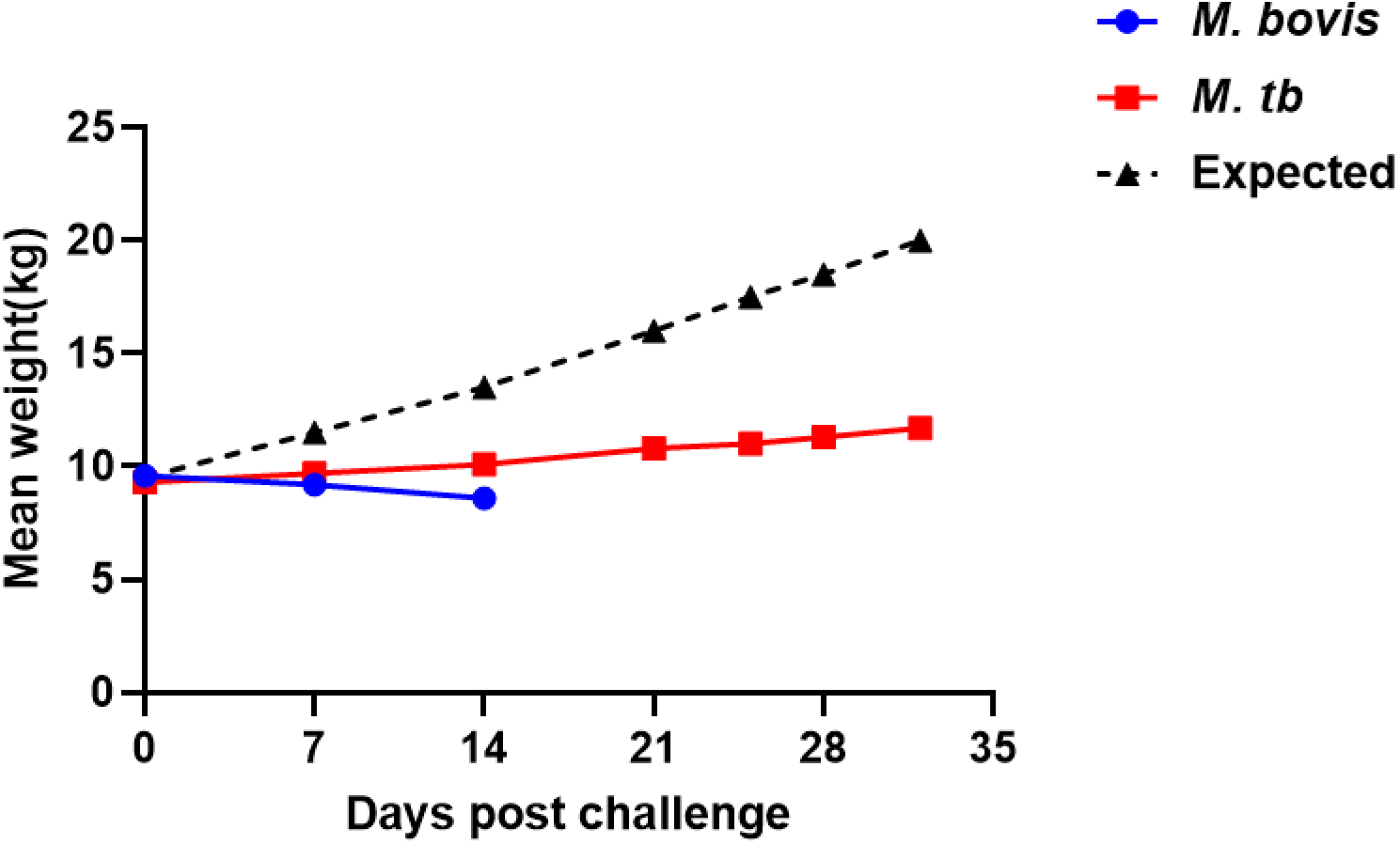
**Mean weight (in Kg) of pigs challenge with *M. bovis* and *M. tb* by intravenous route.** *M. bovis* infected pigs (blue line) started losing weight immediately following the intravenous challenge but the weight measurement could not be taken for the entire trial as the pigs either died or reached their humane endpoint much earlier. In contrast, most *M. tb* infected pigs (red line) gained some weight but it was not close to what is expected for pigs of that breed and age. The expected growth curve (dashed line) is based on data supplied by the Prairie Swine Centre from where the pigs were sourced.

That *M. bovis* IV challenged pigs exhibited rapid onset of morbidity and mortality in this study is consistent with the findings of Bolin *et. al* where pigs IV challenged with 10^8^ CFU of a clinical isolate of *M. bovis* developed severe respiratory distress, fever, exertion and had to be euthanized by day 22 pc [32]. However, Bolin *et. al*. also observed that pigs challenged with a similar dose of *M. bovis* by the intratracheal route exhibited mild symptoms while pigs challenged by tonsillar deposition did not show any signs of infection [32]. This led them to conclude that route of challenge is an important determining factor in the development of overt clinical signs [32]. Our results also highlight MTBC strain is an important determinant of TB disease severity. Additionally, Bolin *et. al.* suggested that the more severe disease seen in their IV challenged pigs was due to the involvement of the central nervous system (CNS) as granulomatous meningitis was found in 3 pigs and *M. bovis* was recovered from the brain tissues and cerebrospinal fluid (CSF) of 4 additional pigs without meningitis [32]. Although we did not sample and analyze the meninges or CSF in our IV challenge trial, Bolin’s findings could explain the higher morbidity and mortality seen with our *M. bovis* challenged pigs. Indeed, studies in other animal TB models have shown high dose bacterial IV challenge can lead to CNS involvement [36, 37].

#### 1.2. Pathology

Detailed necropsy of pigs from both challenge groups revealed the development of edematous, non-collapsing lungs which is indicative of severe inflammation (Figure 4A and B). However, none of the pigs in either challenge groups presented at the gross level with granulomatous lesions on the pleural or visceral surfaces of the lungs (Figure 4A-D). Despite the absence of granulomatous lesions, hemorrhaging on the surface of both the lungs and liver were observed mostly in *M. bovis* challenged pigs (Figure 5A). Moreover, this feature was sometimes accompanied by the presence of fluid in the thoracic and abdominal cavities (Figure 5B).

**Figure 4.**
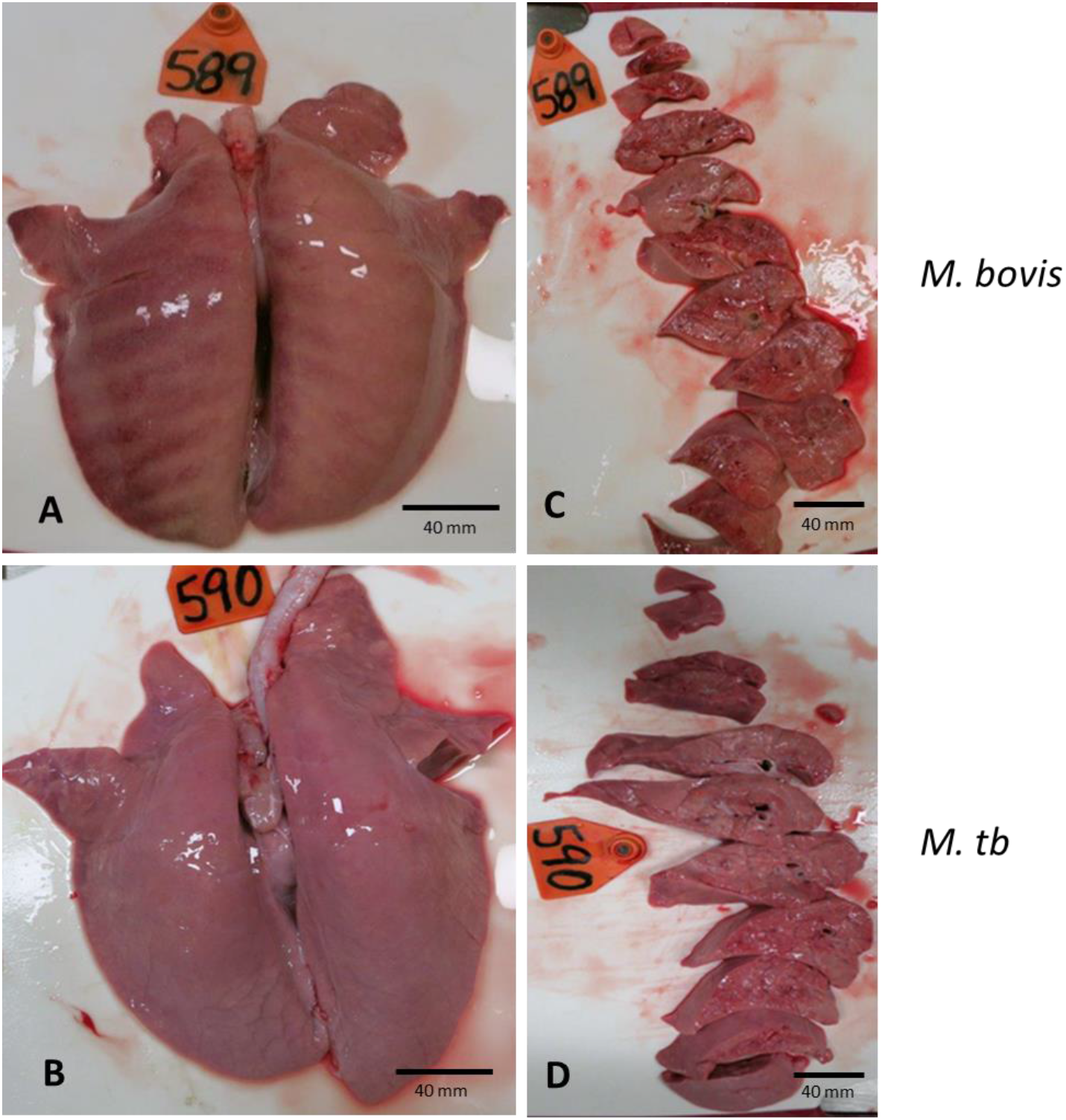
Gross lesions in the lungs of pigs challenged with *M. bovis* and *M. tb* by intravenous route. Lungs from both challenge groups were edematous and non-collapsing but devoid of granulomatous lesions. *M. bovis* infected lung with rib impressions on the dorsal surface (A) and cut section of the same (C). *M. tb* infected lung appears turgid due to edematous inflammation (B) and its cut section (D).

**Figure 5:**
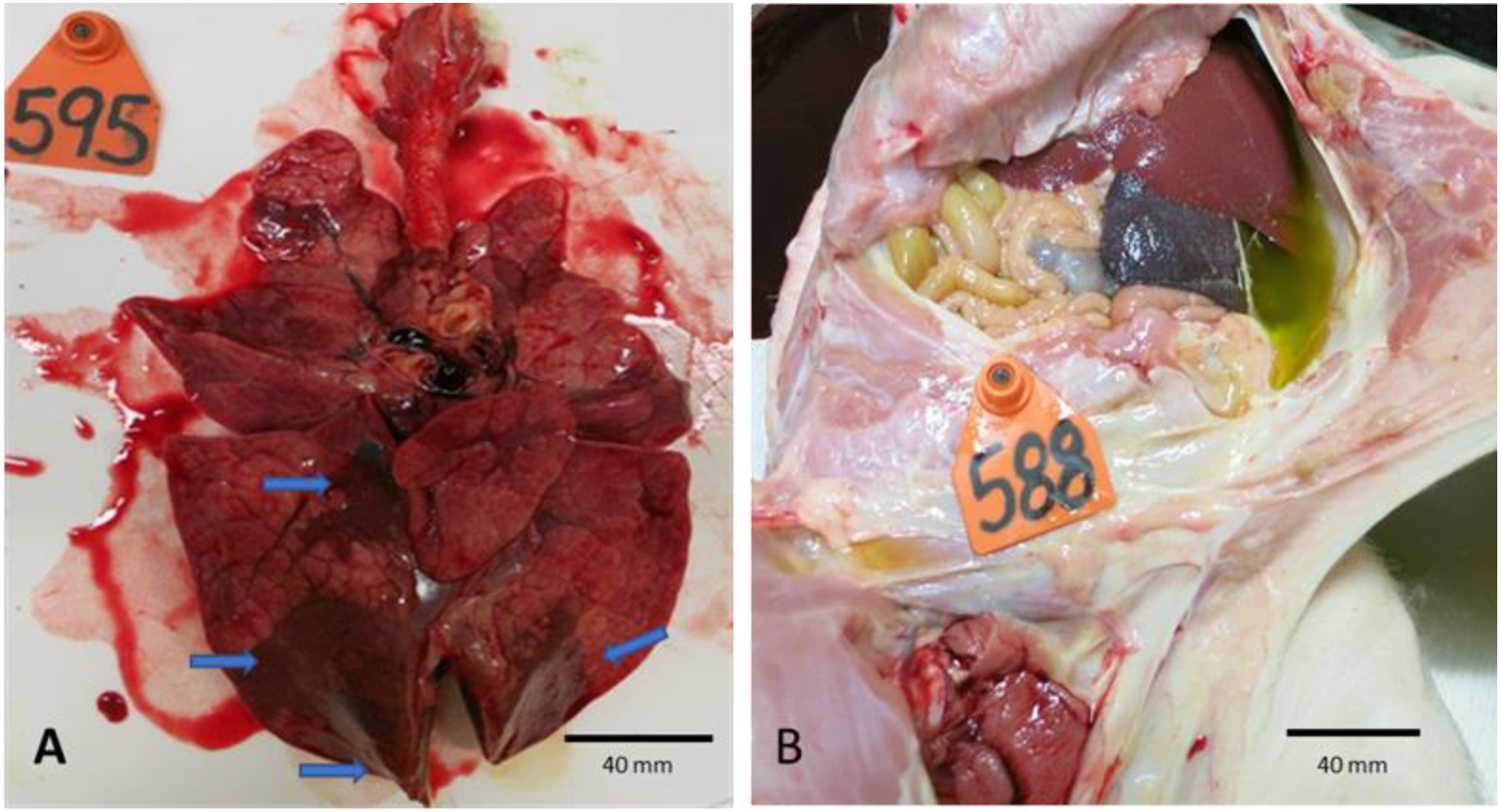
Haemorrhages and effusion associated with *M. bovis* challenged pigs. Some *M. bovis* infected lung had large areas of hemorrhages (indicated by blue arrows) (A). The thoracic and abdominal cavities of some pigs of the same challenge group presented with varying amounts of fluids, possibly due to septicemia (B).

Typical histopathological lesions included loose aggregation of inflammatory cells including macrophages, lymphocytes, and neutrophils (Figure 6A-E). Necrosis with loss of cellular architecture was a common finding in both *M. bovis* and *M. tb* infected animals. Vasculitis and occluding inflammatory cellular aggregates in the arteriole and terminal bronchioles within the lungs were observed (Figure 6A and B). Bronchioles of both *M. tb* and *M. bovis* infected pigs were inflamed with an accumulation of inflammatory cells such as neutrophils, epithelioid macrophages, and multinucleated giant cells (Figure 6C). These occluding lesions are consistent with observations made in previous studies of *M. bovis* infected pigs and cattle [32, 38]. Although areas of necrosis were observed in lung tissues of both *M. tb* and *M. bovis* infected pigs, the extent and frequency of such lesions appeared higher in *M. bovis* infected pigs (Figure 6D). Large areas of necrosis were found in the lymph nodes of *M. bovis* infected pigs while *M. tb* infected pigs tended to display what appears to be small nascent granulomas (Figure 6E). The lesions were microscopic granulomas in early stages not yet surrounded by fibrous capsules. The short duration of the trial most likely did not allow for the development of larger more organized spectrum of granulomas normally seen in naturally infected pigs and humans with active TB. Classification of the granuloma into different stages was not possible in the IV challenge trial because most observed lesions were early inflammatory aggregates without much distinction.

**Figure 6.**
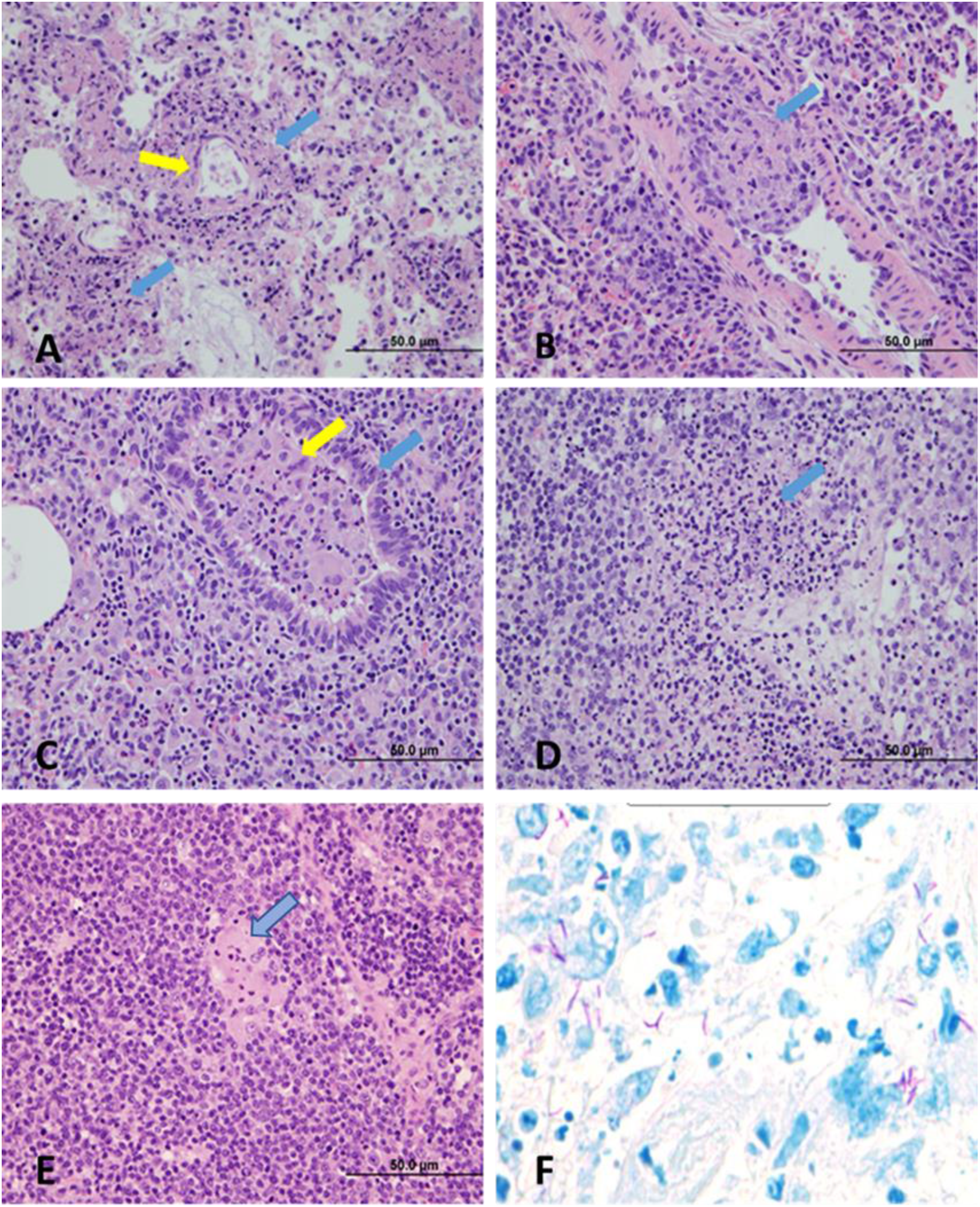
**H&E sections of tissues of pigs challenged with *M. bovis* and *M. tb* intravenously.** *M. bovis* infected lung (40X magnification) with large areas of necrosis (blue arrows) in the alveolar interstitium and the wall of one vessel (yellow arrow) affected by necrosis (A). *M. bovis* infected lung (40X magnification) with an arteriole affected by vasculitis and protrusion of a granuloma (blue arrow) into the lumen (B). *M. tb* infected lung (40X magnification) with a small bronchiole (blue arrow) containing a mixture of neutrophils, epithelioid macrophages, multinucleated giant cells (yellow arrow) (C). *M. bovis* infected lymph node (40X magnification) with large areas of necrosis (blue arrow) and infiltration of neutrophils (D). *M. tb* infected lymph node (40X magnification) with small pyogranuloma (blue arrow) composed of neutrophils and epithelioid macrophages (E). ZN-stained section of *M. bovis* infected cranial lung lobe (F).

#### 1.3. Bacterial burden

Bacteria were recovered from the caudal, mid, and cranial lung lobes, the tracheobronchial lymph nodes, and spleens of both groups of IV infected pigs. Based on the bacterial colony-forming unit (CFU) counts, the bacterial burden was much higher in pigs challenged with *M. bovis* compared to the pigs challenged with *M. tb* (Figure 7). *M. bovis* challenged pigs had on average 10-fold higher bacterial burden in all tissues compared to *M. tb* infected pigs despite being sampled earlier. The difference was also observed in Ziehl-Neelsen (ZN) stained tissue sections from lungs and lymph nodes of the infected pigs (Figure 6F). However, as the route of the challenge was IV, it was difficult to determine whether the presence of bacteria in the spleen and tonsils was due to primary bacterial inoculation or secondary dissemination from the lung. Nevertheless, the lower bacterial burden in *M. tb* challenged pigs suggests *M. tb* but not *M. bovis* replication may be better controlled by pigs.

**Figure 7.**
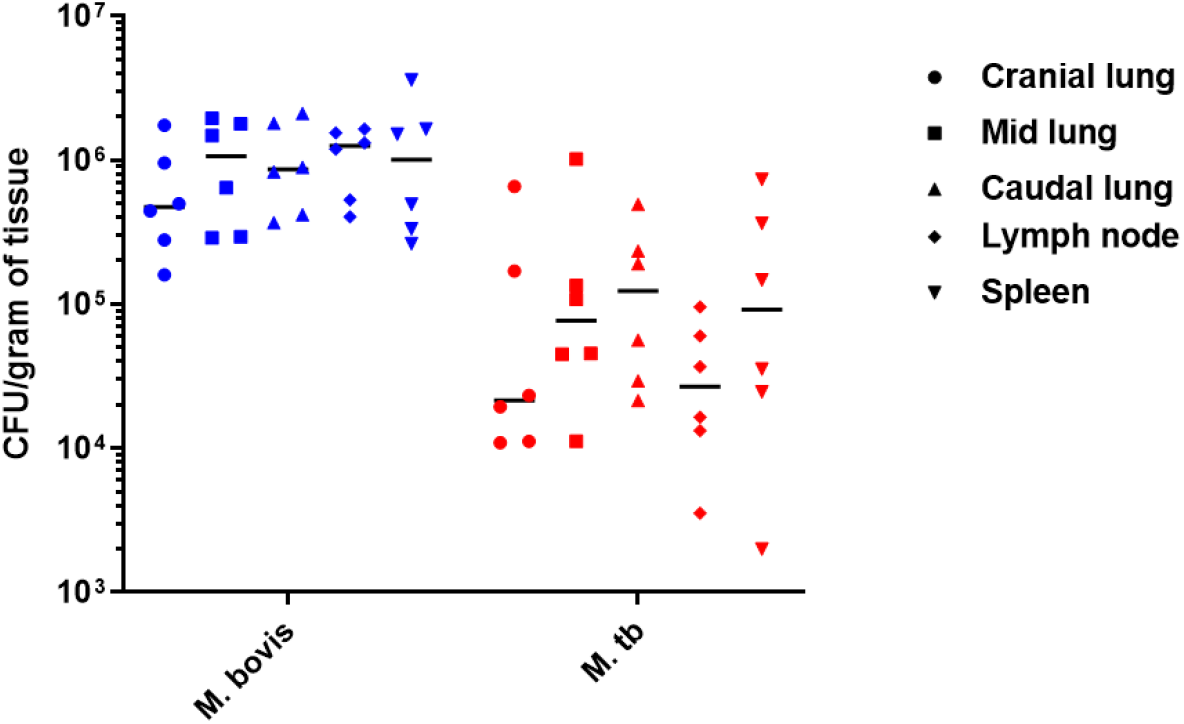
Bacterial burden in *M. bovis* and *M. tb* infected pigs. Each data point represents average CFU per gram of a particular type of tissue in individual animal. Black horizontal line represents median.

Despite being IV challenged with similar doses, the greater morbidity and higher rate of mortality accompanied by marked differences in lack of weight gain, onset of fever, pathology, and bacterial burden in *M. bovis* challenged pigs suggests *M. bovis* is more virulent in the pig model than the human adapted *M. tb*. However, neither *M. tb* nor *M. bovis* IV challenged pigs develop granulomatous lesions due to the short duration of the trial. Also an IV challenge is unlike what occurs during aerosol infection where the disease establishes first as a localized pulmonary infection followed by dissemination to other organs months or years later [39–41].

### 2. Aerosol challenge

#### 2.1. Clinical Signs

Pigs challenged with *M. bovis* and *M. tb* by the aerosol route did not develop overt clinical signs such as cough, fever, and respiratory distress even after 9 weeks post infection. This is consistent with published findings where aerosol infection of minipigs with 10^4^ CFU of *M. tb* failed to generate coughing and fever even after 36 weeks pc [33]. However, significant differences with respect to weight gained by *M. bovis* versus *M. tb* challenged pigs were observed 9 weeks post infection (Figure 8). Pigs challenged with high dose *M. tb* (*M. tb-*HD) registered average weight gains of 44 kg compared to 36 kg in pigs challenged similarly with high dose *M. bovis* (*M. bovis*-HD). However, pigs challenged with low dose of *M. bovis* and *M. tb* (*M. bovis*-LD and *M. tb-*LD) had weight gains of 45 kg and 47 kg respectively (Figure 8). It should be noted that in studies of aerogenic TB in NHPs it takes more than 24 weeks to determine whether an animal has developed latent infection or active disease, based on the presence or absence of clinical signs such as cough and weight loss, AFB in the broncho-alveolar and gastric aspirates, and elevated ESR in the latter [8]. As such, the lack of observable clinical signs in aerosol challenged pigs could be due to the shorter duration of our trial.

**Figure 8.**
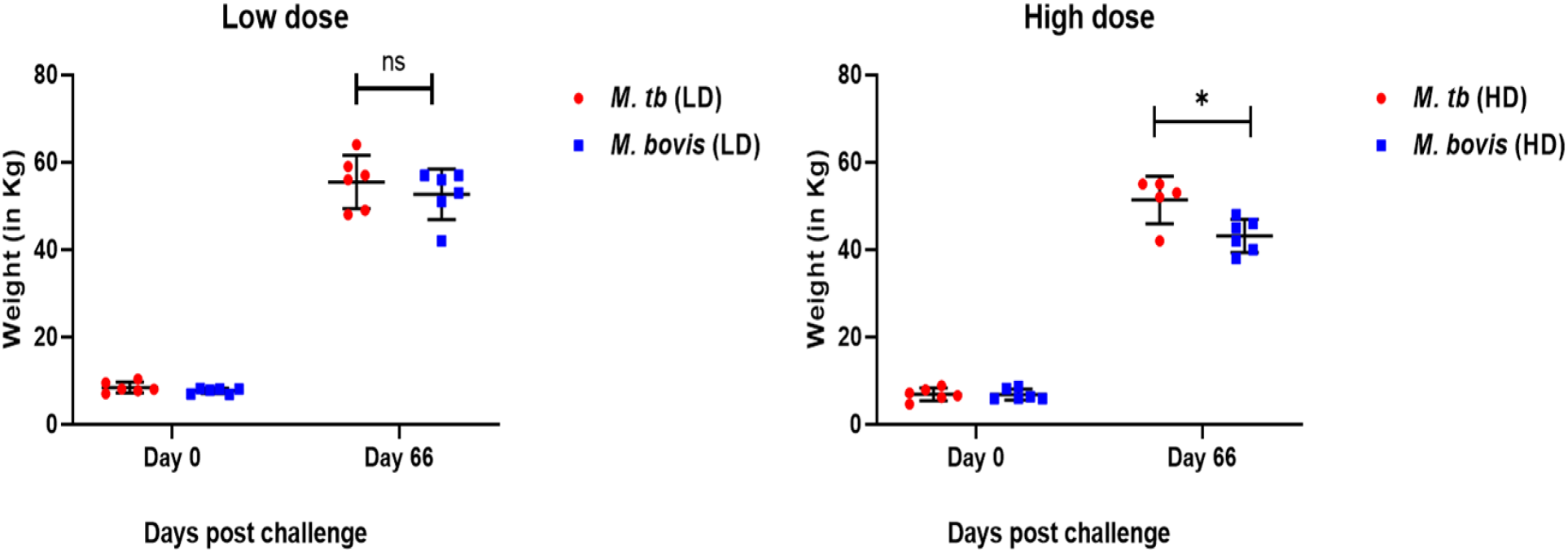
Differential weight gains in pigs aerosol challenged with *M. bovis* and *M. tb*. The weight gained by high dose *M. tb* infected pigs is significantly higher by approximately 8-kg than by high dose *M. bovis* infected pigs (Parametric 2-independent sample T-test: T = 2.961, df = 9, p = 0.016). No significant difference in weight gain was seen in pigs infected with low doses of *M. bovis* and *M. tb* (Each data point represents the actual weight of an individual animal).

#### 2.2. Pathology

Most pigs aerosol challenged with *M. bovis-*HD and *M. tb-*HD exhibited visible pulmonary granulomatous lesions at necropsy 9 weeks post infection. These were uniformly distributed in all the lobes (Figures 9A-D). Entire lungs from pigs of both challenge groups, cut into several slices along their cross-sections further revealed multiple granulomas in the interior surface (Figure 9B and D). Lesions in the lungs varied from very small pin-point hemorrhages to large, calcified nodules with sizes ranging from 2 mm to more than 5 cm in diameter. Notably, cut sections of lungs from *M. bovis*-HD infected pigs presented with large caseating granulomas (Figure 9B). The texture of some of these were hard and gritty due to fibrosis and calcification. Moreover, these mature granulomas were often located close to or encircling the conducting airways (Figure 9B). In several pigs, bronchus stricture was apparent with their walls being affected by caseation indicating possible shedding of bacteria (Figure 9B). In contrast, lungs of pigs infected with *M. tb* showed considerably fewer granulomas (Figure 9C). They were also smaller in size and compact (Figure 9D).

**Figure 9.**
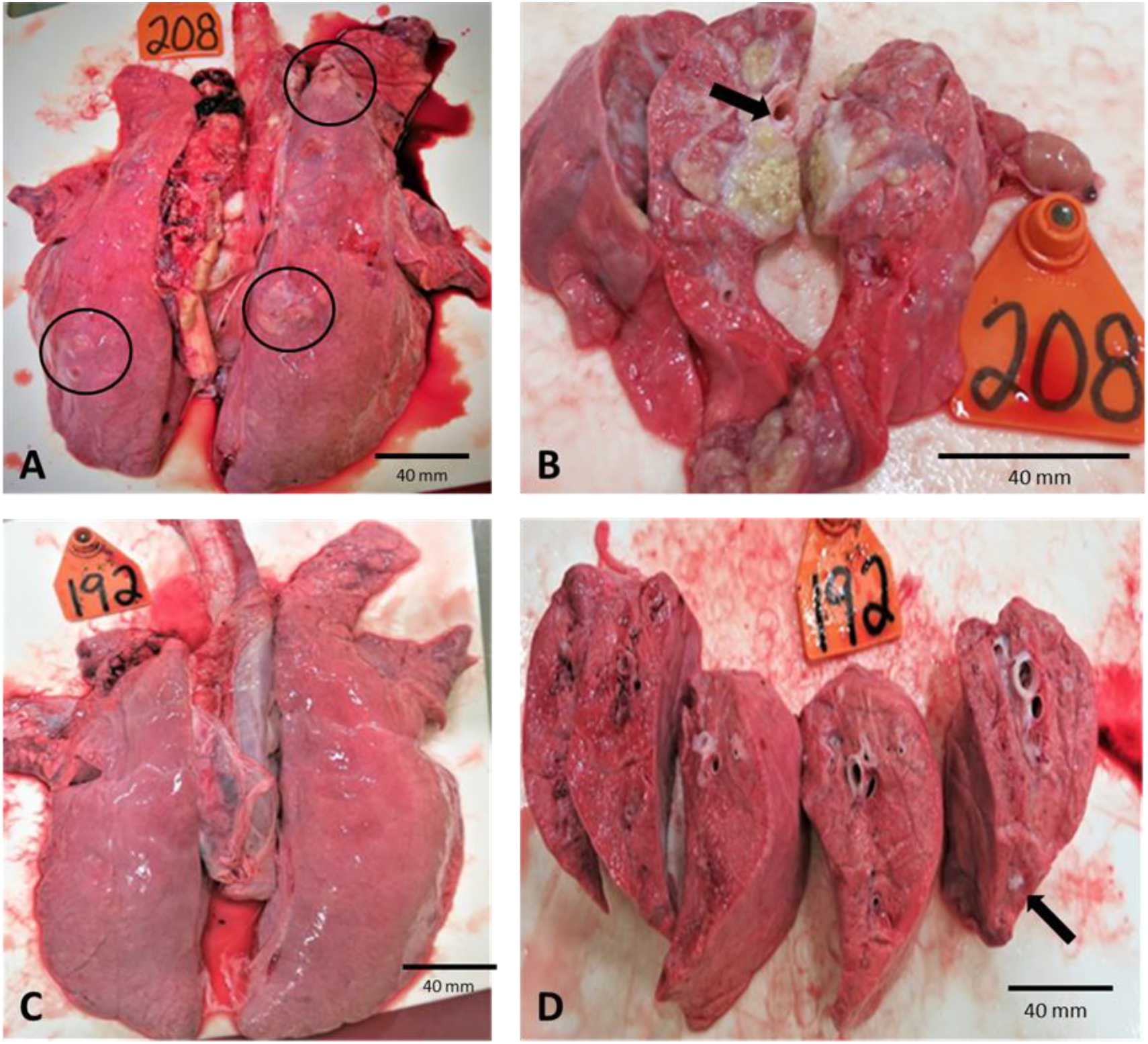
Gross pathology of lungs of *M. bovis* and *M. tb* infected pigs 9 weeks post-challenge. Visible granulomas (in circles) on the surface of the lung in *M. bovis*-HD infected pigs some of which were more than 50 mm in diameter (A). Cut section of *M. bovis* infected lungs with a bronchus surrounded by caseating granulomas (black arrow) (B). *M. tb* infected lung with very few granulomas visible on the surface (C). Cut section of *M. tb* infected lung with few compact, non-necrotizing granulomas in lung parenchyma (black arrows) (D).

Scoring revealed significantly higher lung lesion scores in *M. bovis-*HD infected pigs than both *M. bovis-*LD and *M. tb-*LD infected pigs (Figure 10). Meanwhile, *M. bovis-*HD infected pigs had qualitatively higher lung lesion scores than *M. tb-*HD infected animals, but this was not statistically significant (Figure 10). Likewise, a qualitatively higher but non-significant lung lesion score was observed between *M. bovis-*LD and *M. tb-*LD pigs. Multiple granulomas were observed in the mediastinal lymph nodes (LNs), tracheobronchial LNs and tonsils of all the pigs, irrespective of the challenge strain and dose (Figure 11A-D). Although the granulomas in the LNs were not scored, *M. bovis-*HD infected pigs appeared to have larger and more distinct lesions (Figure 11A). The lesions appeared as large, pale indurations on the serous surface which revealed white caseous masses when cut open (Figure 11A-D).

**Figure 10.**
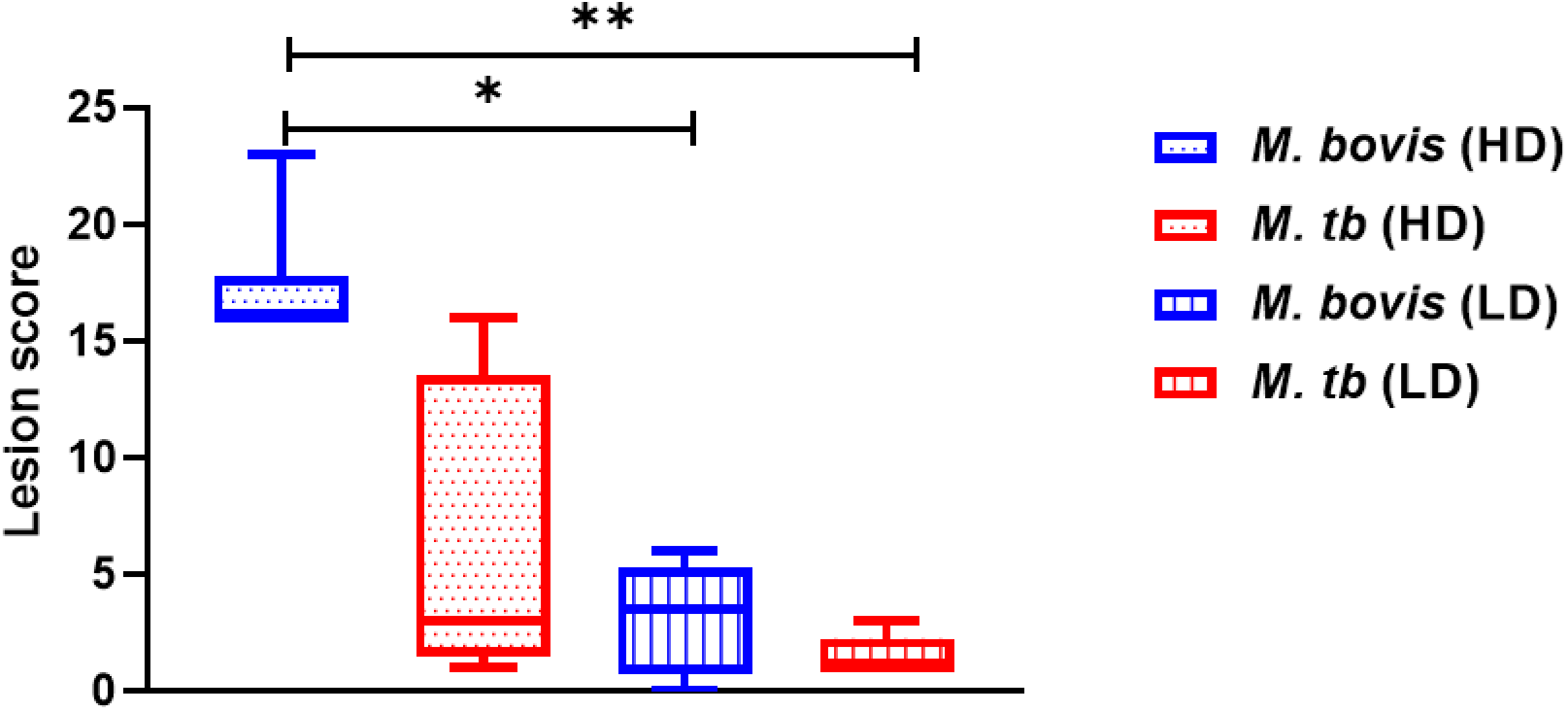
Lung lesion scores of *M. tb* and *M. bovis* aerosol challenged pigs at necropsy 9 weeks post-challenge. Asterisk (*) represents statistically significant difference at 0.05 level of significance and double asterisk (**) represents significance at 0.005 level of significance. Lung lesion scores between different treatment groups were compared by non-parametric Kruskal-Wallis Test (α = 0.05). Significant differences in lung lesion scores were found based on treatment groups (Kruskal Wallis H = 14.016, df = 3, p = 0.003). Subsequently, multiple pairwise comparison using Mann-Whitney U test was done to determine statistical significance between the treatment groups. Lung lesion scores in *M. bovis*-HD was significantly higher than *M. bovis*-LD (Mann Whitney U = 10.33, p = 0.04) as well as *M. tb*-LD (Mann Whitney U = 13.83, p = 0.002). Any other pairwise combinations were non-significant. P value reported have been adjusted with Bonferroni correction for multiple tests. *M. bovis*-HD infected pigs also had qualitatively more severe lung pathology compared to *M. tb-*HD infected pigs. There was no significant difference between the pigs infected with low doses of *M. bovis* and *M. tb*.

**Figure 11.**
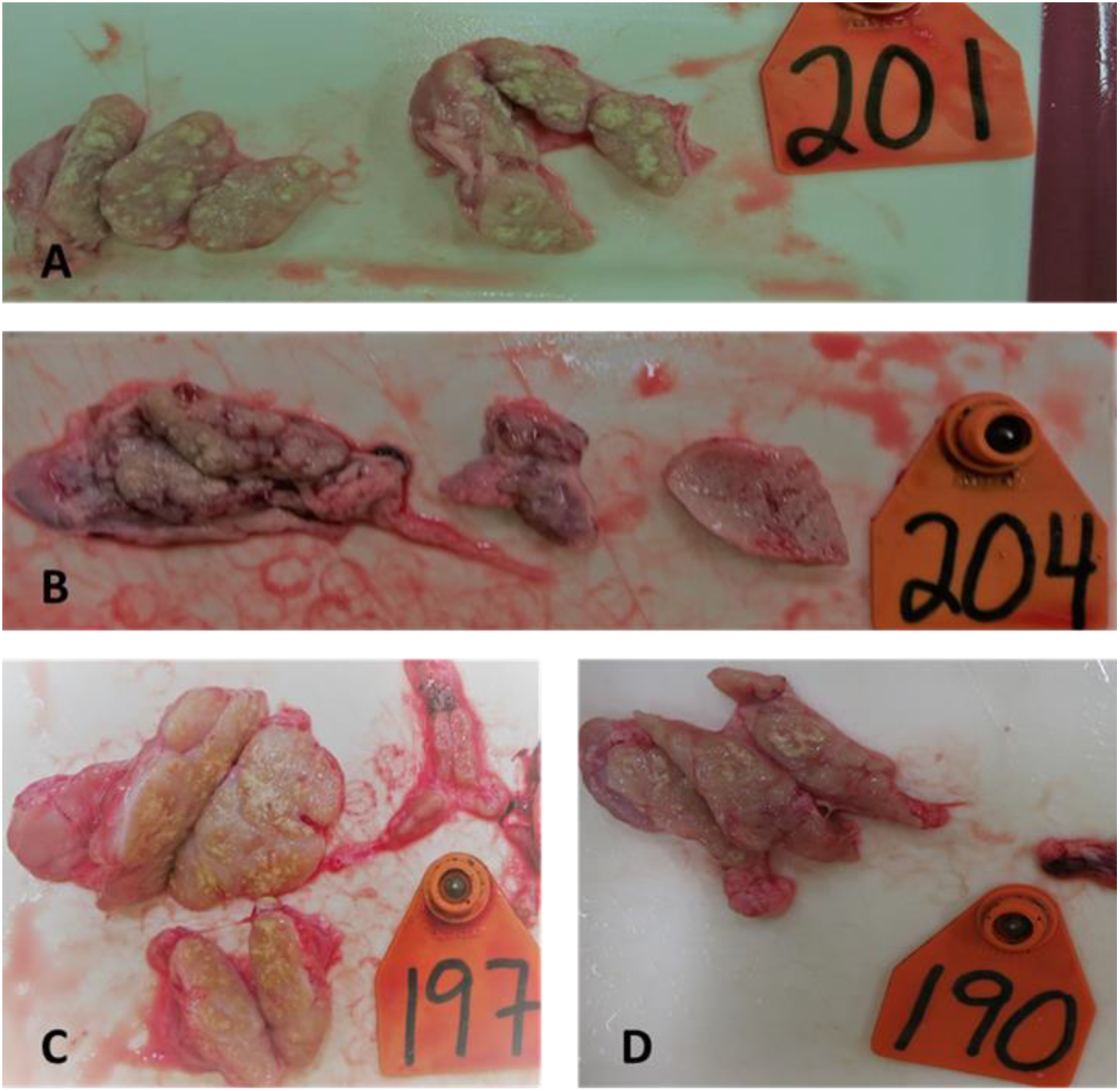
Gross pathological lesion in lymph nodes of *M. bovis* and *M. tb* aerosol infected pigs. Cut sections of tracheobronchial and mediastinal lymph nodes from *M. bovis*-HD infected (A), *M. tb*-HD infected pigs (B), *M. bovis*-LD infected (C) and *M. tb*-LD infected pigs (D). Note whitish yellow caseous mass in the cut sections of lymph nodes. Lesions are more abundant in *M. bovis-*HD infected pigs which corelates with higher bacterial burdens in lymph nodes of these animals.

Microscopically, pigs aerosol challenged with either *M. tb* or *M. bovis* showed granulomatous lesions in at least one organ (Figure 12A-F). These include lesions with thickening of lung parenchyma (Figure 12A), multiple satellite granulomas (Figure 12B), collapsed bronchioles (Figure 12C), and with accumulations of macrophages, lymphocytes, and neutrophils (Figure 12D), and organized granulomatous lesions surrounded by thick fibrous capsule (Figure 12D). Advanced stage lesions with fibrosis encapsulating granulomas (Figure 12E) as well as with necrotic cores (Figure 12F) were also observed. We also observed foamy macrophages, epithelioid macrophages and multinucleated giant cells that comprise the core of granulomas (Figure 13). A large number of neutrophilic infiltrations was observed in advanced stage granulomas indicating the start of liquefactive necrosis (Figure 13). ZN staining showed varying numbers of acid-fast bacilli (AFB), which did not appear to be associated with CFU burden in our analysis. Interestingly, Santos *et. al.* reported that AFB cannot be reliably associated with the presence or absence of infection in wild pigs [42]. Moreover, there were differences in the types of granulomas observed even within the same pig, a feature commonly associated with active TB in cynomolgus macaques [43]. Due to the differences in the number and nature of granuloma in different challenge groups in this study, a classification scheme was developed.

**Figure 12.**
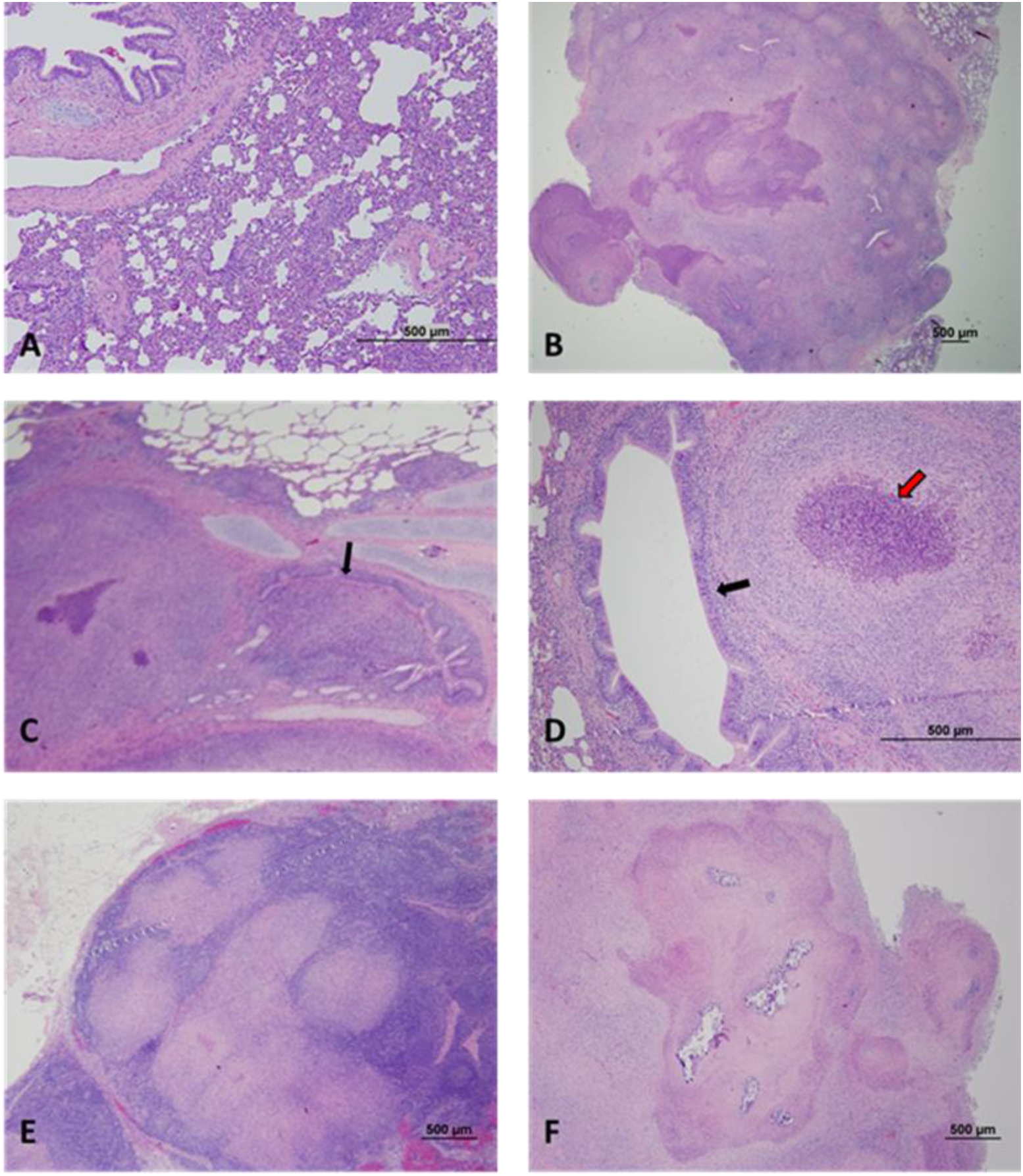
Histopathology of pigs infected with *M. bovis* and *M. tb*. Thickening of lung parenchyma can be seen in *M. tb*-LD infected lung (A). Multiple satellite granulomas with a follicular appearance surrounding central necrotic core of *M. bovis*-HD infected pig lung (B). *M. bovis*-HD infected lung with advanced stage granuloma with collapsed bronchiolar lumen and infiltration (arrow) (C). Another *M. bovis*-HD infected lung with stricture of bronchiole by a large granuloma (black arrow) and infiltration of neutrophils (red arrow) suggesting beginning of necrotic process (D). *M. tb*-HD infected mediastinal lymph node with a multifocal lesion and thick capsule surrounding those lesions (E). *M. bovis*-HD mediastinal lymph node with advanced stage lesion, amorphous mantle and liquefying core (F).

**Figure 13.**
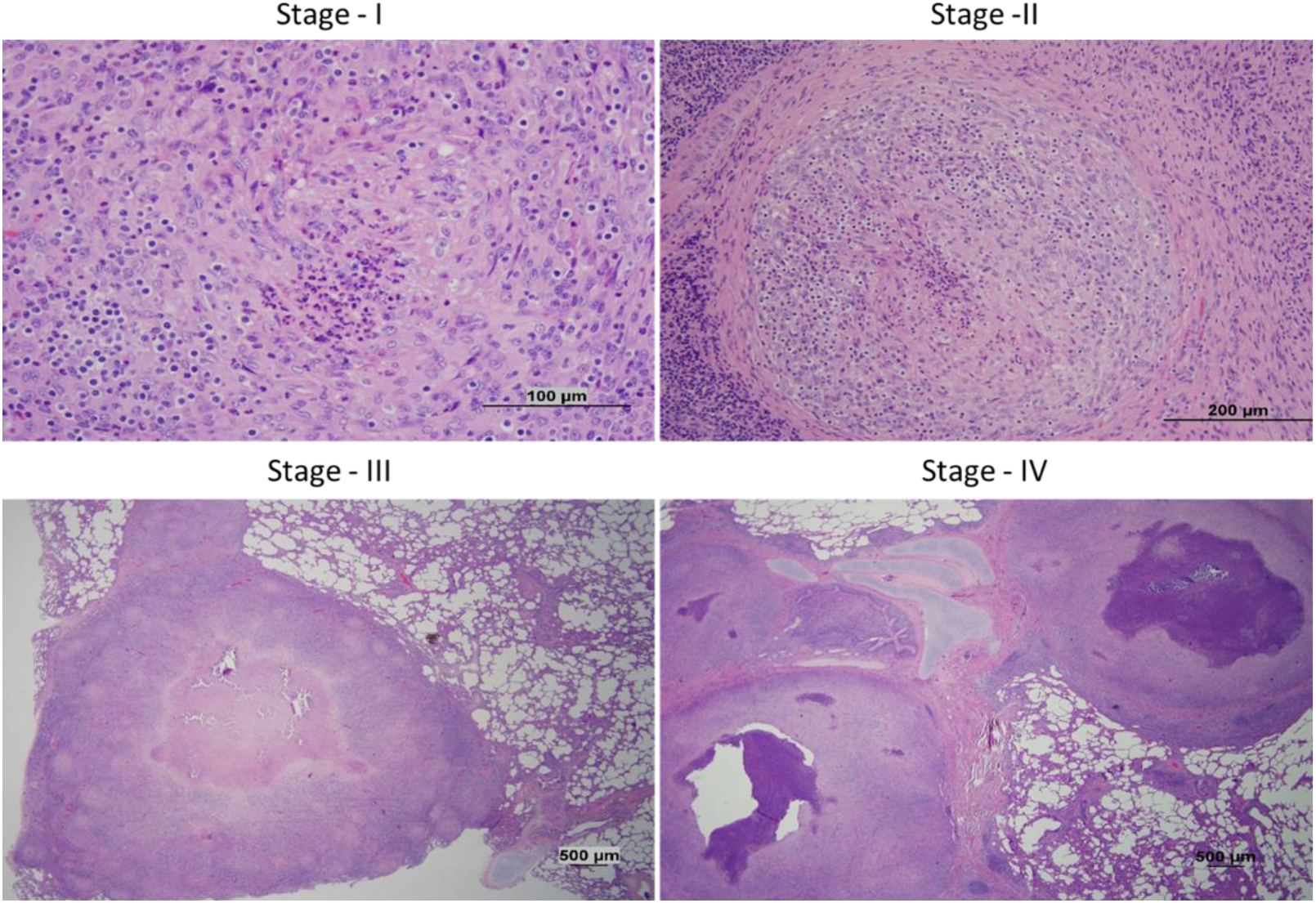
Different stages of granuloma in pigs infected with *M. bovis* & *M. tb*. At least one lung H&E-stained sample from all the animals in each group were studied and the granulomas were categorized into one of the four stages. Stage-1 lesions are the initial aggregation of inflammatory cells such as epithelioid macrophages, lymphocytes, neutrophils. There is a lack of fibrous capsule, and the structure is unorganized. Stage-2 granulomas appear as organized structure with a central core of neutrophils, foamy macrophages and abundant epithelioid macrophages, and a peripheral rim of T and B lymphocytes. A fibrous capsule demarcates the granuloma from surrounding parenchyma. The difference between stage-2 and stage-3 granulomas is the formation of necrotic centers within the granulomas and thickening of the capsule. Stage-4 granuloma have extensive caseous necrosis and liquefaction leading to an amorphous zone in the center of the granuloma. Multiple granuloma may coalesce to form a large, multifocal lesion as shown in the bottom-right panel.

Histologically, lung granulomas can be classified into different stages [31,44,45]. Stage-1 granulomas are characterized by the loose aggregation of inflammatory cells such as epithelioid macrophages, lymphocytes, neutrophils, and very few multinucleated giant cells (MNGCs) without fibrous capsule [31]. Stage-2 granulomas are characterized by fibrous encapsulation of an organized lesion composed of the central core of neutrophils, foamy macrophages and abundant epithelioid macrophages, and a peripheral rim of T and B lymphocytes [31]. Stage-3 granulomas have much thicker and complete fibrous encapsulation and while the cellular constituents are similar to stage-2, there is more neutrophilic infiltration towards the core of the lesion which ensues necrosis [31,44,45]. Smaller granulomas may begin to fuse to form a multifocal granuloma at this stage. Stage-4 granuloma is marked by extensive caseous necrosis and liquefaction such that there is an amorphous to acellular zone in the center of the granuloma. This is when the granuloma loses its integrity and the animal is believed to be transmissive [46]. Stage-4 lesions may be surrounded by few to numerous satellite granulomas in stage-1 or stage-2 [31].

In our study, at least one lung H&E-stained sample from all the animals in each group of the aerosol challenge trial was studied and the granulomas were categorized into one of the four stages described above (Figure 13). For analysis, stages-1 and 2 lesions were considered as early and intermediate granulomas respectively whereas stages-3 and 4 were considered as the advanced stage granulomas. It was interesting to note that 9 (81%) out of 11 lung samples from *M. bovis-* HD infected pigs had advanced granuloma suggesting that these animals are manifesting an active form of TB (Table 2). In contrast, only 3 (50%) out of 6 lung samples had an advanced lesion in *M. tb*-HD infected pigs (Table 2). Similarly, only 1 out of 11 tissues had stage-4 granuloma in *M. bovis*-LD challenged pigs while no stage-4 granulomas were found in *M. tb*-LD infected pigs (Table 2).

In this study, we have provided a detailed characterization and classification of histopathological lesion in pigs which is an advancement over the existing body of knowledge available from previous works on minipigs [29, 33]. Although the number of tissue samples available from the aerosol challenge trial for H&E staining were few, there is clear indication that infection with *M. bovis*-HD results in the production of more advanced stage granulomas than infection with *M. tb-*HD. Significantly, this demonstrates pigs can develop caseous granulomas with central necrosis – a hallmark of active human TB [47]. A comparative study done in cattle reported that *M. bovis* challenged calves had lesions ranging from stage 1 to 4 while the calves challenged with two different strains of *M. tb* had arrested granuloma development with lesions mostly at stages 1 and 2 [34]. The differences in pathological outcomes observed in our study is consistent with the cattle study, which attributed the difference in pathology to attenuation of *M. tb* in the animal host [34, 48]. This finding suggests that *M. tb* could be used to generate latent infection model while *M. bovis* could be used to develop an active disease model in pigs, although the dose might have to be adjusted to a fewer CFU per animal to mimic latent infection in humans.

#### 2.3. Bacterial burden

Bacteria were only recovered from the lungs (pooled data of cranial, mid, and caudal lobes tissues) of 2 out of 6 pigs for both *M. bovis-*LD and *M. tb-*LD challenged groups (Figure 14). Similar bacterial burden was seen for lymph nodes (pooled data of tracheobronchial, mediastinal, and retropharyngeal lymph node tissues) of *M. bovis-*LD and *M*. *tb*-LD challenged groups (Figure 14). Bacteria was not recovered from the spleens of either *M. tb*-LD or *M. bovis-*LD challenged pigs (Figure 14). With the HD challenge groups, bacteria were recovered from the lungs (pooled data of cranial, mid, and caudal lobes tissues) of *M. tb*-HD challenged pigs and *M. bovis-*HD challenged pigs. Notably, lungs samples of *M. bovis-*HD challenged pigs had higher median burdens. Similarly, there was a higher bacterial burden in the lymph nodes (pooled data of tracheobronchial, mediastinal, and retropharyngeal lymph node tissues) of *M. bovis-*HD challenged pigs compared to *M. tb*-HD challenged pigs (Figure 14). Viable bacteria were recovered from the spleens of two pigs challenged with *M. bovis-*HD only, suggesting post-primary dissemination likely through hematogenous route. A similar finding was reported in goats where dissemination of infection outside the thorax was seen only in *M. bovis* infected goats [35]. Overall, pigs aerosol challenged with *M. bovis* were found to have a relatively higher pulmonary burden than *M. tb*, especially in the high dose challenge groups. This suggests that pigs are able to contain infection with *M. tb* better than infection with *M. bovis*. Indeed, mycobacteria were recovered from the lungs of only 3 *M. tb* challenged pigs. This was despite the similar challenge dose and duration of infection for *M. tb* and *M. bovis*. The differences in tissue bacterial burden also correlates positively with the difference in the lung lesion scores. Our study also demonstrates heterogeneity in bacterial burden and lesion severity which has previously been reported in the minipig model as well as in cattle [33, 49]. Furthermore, the differences in pathology and bacterial burden are likely associated with the differences in overall weight gains exhibited by *M. bovis*-HD and *M. tb* -HD infected pigs.

**Figure 14.**
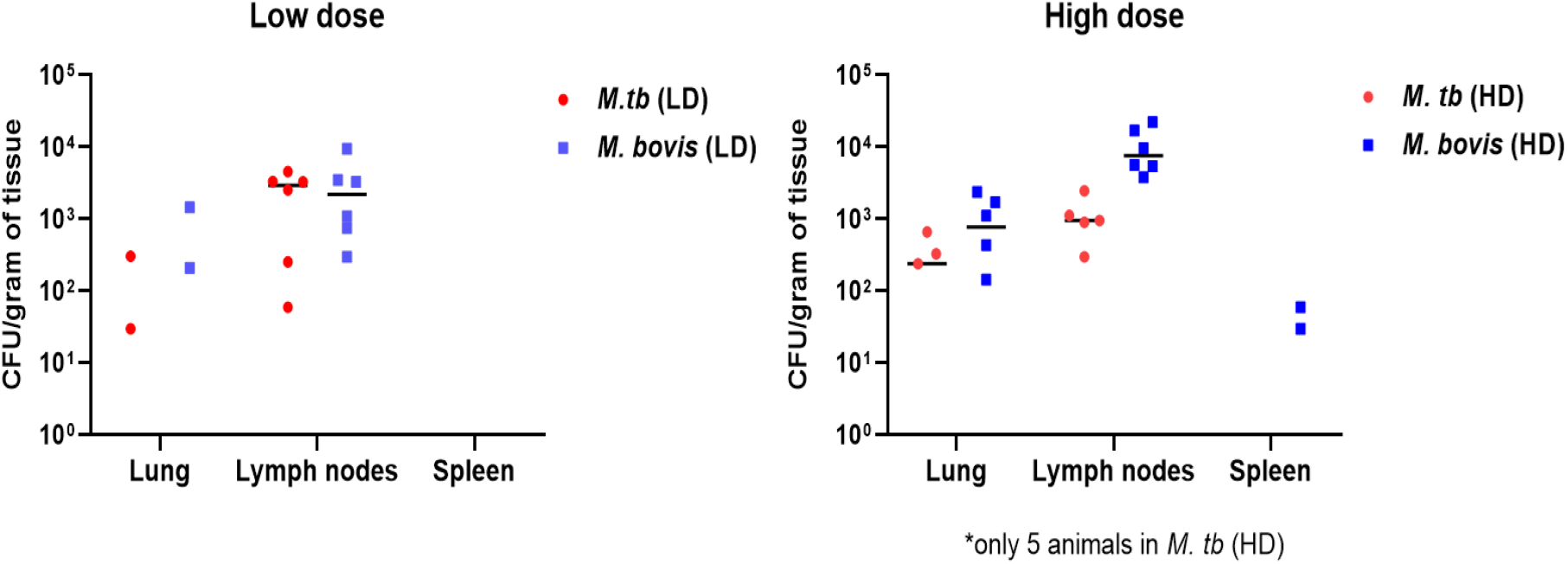
**Bacterial burden (CFU per gram of tissue) in different organs of pigs infected with high and low doses of *M. bovis* & *M. tb***. Each point in the graph represents CFU/g of one animal and the black line represents the group median. Value for some individuals is missing as it was below the detection limit. Recovery of mycobacteria from spleen samples of two *M. bovis* HD challenged pigs indicates post-primary dissemination.

#### 2.4. Peripheral T-cell response after challenge with aerosolized *M. bovis* and *M. tb*

TB disease progression in aerosol challenged pigs was done using the Interferon-gamma release assay (IGRA), a sensitive and specific test for diagnosing TB in pigs [50]. The assay quantifies the production of Interferon gamma (IFN-γ) in response to mycobacterial antigens by infected animals. A steady increase in IFN-γ concentrations over time was observed in all four treatment groups irrespective of the doses and strains used (Figure 15). The immune conversion of IGRA from negative to positive typically occurs within 4-6 weeks post infection in humans [51]. In this study, most pigs showed detectable IFN-γ levels 4 weeks after challenge which correlates with the beginning of the adaptive immune response and activation of effector T-cells, which are the main producers of IFN-γ. The concentration reached a maximum value at 9 weeks post-challenge suggesting the highest T-cell immune activity at that period. Similar kinetics of T-cell immune response is documented in minipigs [29], wild pigs [52], goats [53] and cattle [34]. Moreover, infection of pigs with *M. tb* and *M. bovis* at both high and low doses induced similar peripheral immune responses. This is consistent with findings from a study comparing IFN-γ responses to bPPD in cattle that were also infected with *M. bovis* and *M. tb* for 16 weeks but displayed no differences in IFN-γ responses [54].

**Figure 15.**
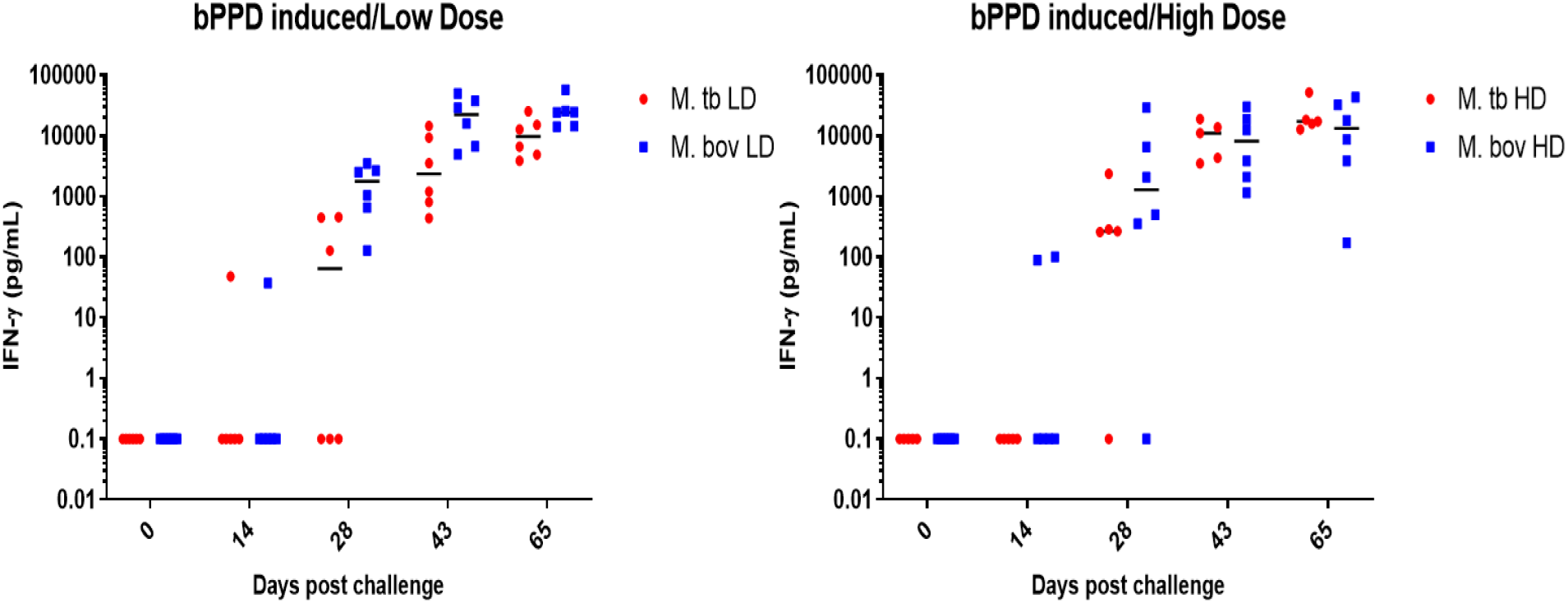
IFN-γ response due to bPPD stimulation of the whole blood. Each data point indicates concentration of IFN-γ in individual animal at that point after removing background concentration (PBS stimulated sample). Although few pigs had measurable IFN-γ response by day 14 post challenge, most pigs displayed IFN-γ response at day 28 post challenge. Despite challenge with different doses, the peripheral IFN-γ response is similar for *M. bovis* and *M. tb* challenge.

## Conclusion

In this study, we have determined that *M. bovis* AF2122/97 is more virulent in pigs compared to *M. tb* Erdman regardless of route of infection. Our extensive analysis of the histopathological lesions found in the tissues of both *M. bovis* and *M. tb* infected pigs revealed domestic pigs mimic the pathological features of a human TB. Specifically, pigs aerosol-challenged with high doses of *bovis* developed wide spectrum of granulomatous lesions including caseation and necrosis, much like in humans with active TB, while pigs challenged with low dose *M. tb* exhibit arrested granuloma development much like in humans with latent TB. These findings support the notion that the domestic pig is a useful animal model to study active and latent TB.

## Materials and Methods

### 1. Preparation of challenge material

Frozen glycerol stock of *Mycobacterium tuberculosis* (Erdman) and *M. bovis* (AF2122/97) were thawed and plated on Middlebrook 7H11 agar plates. A single colony was used to inoculate Middlebrook 7H9 liquid medium, supplemented with 10% Albumin Dextrose Catalase (ADC) and 0.05% Tween-80. *M. bovis* culture media was additionally supplemented with 0.5% sodium pyruvate. Bacterial cultures were incubated at 37^0^C until mid-log phase growth was achieved. A subculture was prepared with a starting OD_600_ of 0.1 twenty-four hours before the animal challenge. On the day of challenge, bacterial culture was spun at 300g for 3 minutes and supernatant, which contains single-cell suspension, was harvested for enumeration by OD. An OD of 1 corresponds to 3 x 10^8^ CFU/mL for *M. tb* and 1 x 10^8^ CFU/mL for *M. bovis.* Necessary volumes of each culture were diluted in PBS to make the required challenge inoculum.

### 2. Study Design and Challenge

#### • Intravenous challenge

Twelve 4-weeks old, domestic pigs were sourced from Prairie Swine Center and brought to Agriculture Biosafety Level-3 (Ag-BSL-3) facility at VIDO. Animals were tagged with unique numbers and randomly divided into two groups (Group A and Group B) with 6 pigs in each group (Table 1). The pigs were allowed to acclimatize in the containment facility for 7 days before the challenge. Group A was challenged with intravenous injection of 3 × 10^8^ CFU of *M. bovis* per pig while group B was challenged with intravenous injection of 3 × 10^8^ CFU of *M. tb* per pig. Six pigs from each challenge group were housed in 14 feet x 5.5 feet pens. They were kept on a raised platform and were under continuous surveillance through a camera and in person. Although both the challenged groups were housed in the same containment room, they were separated by a 2.5 feet high physical barrier enclosing each pen. The animal containment rooms had a net negative pressure with the air exchange maintained at 10 cycles per hour allowing for whole air volume replacement multiple times in an hour. The exhaust mechanism thereby greatly minimizes the chances of cross-infection. The room was provided with 12-12 hours of light and dark cycles. The animals were monitored for changes in clinical signs such as respiratory distress, fever, anorexia, weight loss, and other morbidities. Daily weight and temperature measurements were taken for 10 consecutive days. Food and water were provided ad-libitum for the entire period of the experiment.

**Table: 1.** **Lung lesion scoring scheme for cattle** (developed by Drs. Carly Kanipe & Mitch Palmer, USDA)

| Lesion score | Observation |
| --- | --- |
| 0 | No visible lesion |
| 1 | **1.** Less than 10 lesions<br>**2.** All lesions less than 10 mm in diameter |
| 2 | **1.** 6-10 lesions<br>**2.** Rare ($\leq 2$) lesions between 10-20 mm in diameter |
| 3 | **1.** 11-20 lesions<br>**2.** Occasional (3-5) lesions between 10-20 mm in diameter |
| 4 | **1.** More than 20 lesions<br>**2.** Frequent ($\geq 5$) lesions between 10-20 mm in diameter |
| 5 | **1.** Countless & coalescing lesions or<br>**2.** More than half of the lesion is $>10$ mm or |
|  | **3. Any lesions > 20 mm in diameter** |

**Table 2:** Lung histopathological lesion scoring of pigs infected with M. bovis & M. tb. At least one H&E slide from each animal from all four groups was analyzed. NA -not assessed

| Groups | Animal ID | Cranial Lobe | Middle Lobe | Caudal Lobe |
| --- | --- | --- | --- | --- |
| *M. bovis*<br>HD | 191 | 2 | 4 | NA |
|  | 196 | 4 | NA | 4 |
|  | 200 | 1 | NA | 3 |
|  | 201 | 3 | NA | NA |
|  | 208 | 4 | 4 | NA |
|  | 212 | 4 | 4 | NA |
| *M. bovis*<br>LD | 189 | 1 | NA | 1 |
|  | 193 | 3 | NA | 1 |
|  | 197 | 4 | NA | NA |
|  | 202 | 2 | 1 | NA |
|  | 207 | 1 | NA | 1 |
|  | 209 | 3 | NA | 1 |
| *M. tb*<br>HD | 192 | 4 | NA | NA |
|  | 194 | 2 | 4 | NA |
|  | 199 | 1 | NA | NA |
|  | 204 | 3 | NA | NA |
|  | 206 | 2 | NA | NA |
| *M. tb* | 190 | 3 | NA | NA |
|  | 195 | 1 | 2 | NA |
|  | 198 | 1 | NA | NA |
| LD | 203 | 1 | NA | 1 |
|  | 205 | 1 | NA | NA |
|  | 211 | 2 | 2 | NA |

#### • Aerosol challenge

24 three-week-old domestic pigs were brought to the containment facility at VIDO and allowed to acclimatize for a week. These animals were tagged and randomly divided into four groups containing 6 animals each. Two challenge doses were prepared for each strain as mentioned above and the inoculum was aerosolized through a Collison nebulizer (CH Technologies, USA), connected to a tank of compressed oxygen. Aerosol generated was passed through a tube into a mask that fitted snugly to the pig’s snout. Each pig received 2 mL of the challenge inoculum. Post-challenge, the animals were housed similarly as described for the IV challenge.

### 3. Gross pathology and Histopathology

Pigs that died during the experiment or those euthanized were subjected to a detailed gross and histopathological examination. During necropsy, the presence of tuberculosis-associated lesions was monitored in the lungs, liver, spleen, lymph nodes and other visceral organs in the thoracic and cervical regions. For histopathological evaluation, a 2x2x2 cm^3^ section of tissue was sampled and secured within a tissue cassette, immersed into 10% buffered formalin for fixation inside the BSL-3 containment. These samples were later taken to Prairie Diagnostic Services, Saskatoon for Hematoxylin-Eosin (H&E) and Ziehl-Neelsen (ZN) staining.

#### • Lung lesion scoring

Using visual and palpation skills, the chief veterinarian at VIDO scored each lung lobe based on the scoring scheme as developed by Palmer *et. al.* in cattle with minor modifications [38] (Table 1). The total lung lesion score for each animal was obtained by adding the lesion score of individual lobes. Both sizes of the granuloma and number were considered while scoring the lesions.

#### • Staging of granulomas

Tissue sections were categorized into different stages based on their cellular composition, architecture, degree of necrosis and extent of fibrosis [31,44,45] For each H&E stained tissue sample, 10 field-of-views were chosen randomly within a slide and each field was categorized into one of the four stages. The sample would then be classified into one of the four stages if the occurrence of one particular lesion was in at least 5 field-of-views (50% cutoff limit). Stage I indicates early-stage loose aggregation of cells due to localized immune response whereas stage IV indicates advanced stage granuloma with caseation and liquefactive necrosis. This stage is considered to represent transmissible pathology. Only lung tissue sections were examined for classification.

### 4. Evaluation of tissue bacterial burden

Approximately 1 g of tissue was obtained from at least three different sections of the lungs and lymph nodes using a sterile biopsy punch. The samples were stored in 3 mL of PBSA + tween-80 (0.05%) + ampicillin (80µg/ml) + cycloheximide (100µg/ml) and frozen. Later, the tissue samples Frozen samples were homogenized, and different dilutions were plated in Middlebrook 7H11 selective media. Tissue homogenizer was washed thoroughly to prevent carry over of bacteria from one sample to the other. Once grown, the colonies were enumerated using automated colony counter (Scan Interscience).

### 5. Interferon-gamma release assay (IGRA) protocol

The progression of the disease was monitored by a blood-based interferon-gamma release assay. Blood samples from each pig were collected a day before the challenge and every two weeks post challenge. 10 ml blood per animal was collected in lithium heparin tubes, placed in a rocker to allow proper mixing with an anticoagulant. A 24-well tissue culture (Costar, Corning Incorporated, USA) plate was set up with 100 µL of mycobacterial antigen per well. The antigens used were bovine PPD (300 IU/mL), EsxA (5 µg/mL) and PBS control. Mycobacterial culture filtrate contains soluble antigens secreted by the bacteria in the culture. 1.5 ml blood from each animal was transferred to individual wells and mixed by gentle rotation of the plates. The plate was covered with a sterile breathable membrane and incubate at 37^0^C for 24 hours. The next day, the breathable membrane cover was removed, and the plate was spun at 800 g for 15 mins at room temperature. Plasma was harvested from the wells without disturbing the cell pellet and transferred into Spin-X centrifuge tube filter (Corning Incorporated, USA) and spun at 15000 g for 20 mins at room temperature. The filtrate obtained was transferred to a fresh screw-capped tube and stored at -80^0^C until used.

### 6. Statistical Analysis

The rate of survival between IV challenged *M. bovis* and *M. tb* was compared by Log-rank test. Comparison of weight gained by pigs in different groups in the aerosol challenge experiment was done by 2-independent sample T-test. Likewise, comparison of lung lesion scores was done by Kruskal-Wallis test with subsequent multiple pairwise comparisons using Mann-Whitney U test.

## Acknowledgements

VIDO receives operational funding from the Government of Saskatchewan through Innovation Saskatchewan and the Ministry of Agriculture and from the Canada Foundation for Innovation through the Major Science Initiatives for its CL3 facility. The research was supported by grants awarded by the Saskatchewan Health Research Foundation and the National Sanatorium Association of Canada to JMC. NN, KD are recipients of scholarships from the department of Veterinary Microbiology, WCVM. The authors would like to acknowledge VIDO Animal Care team for their help in conducting the animal trials. The article is published as VIDO manuscript series no. 956.

## Notes

### Competing Interest Statement

The authors have declared no competing interest.

